# An integrated strain-level analytic pipeline utilizing longitudinal metagenomic data

**DOI:** 10.1101/2022.02.15.480548

**Authors:** Boyan Zhou, Chan Wang, Gregory Putzel, Jiyuan Hu, Menghan Liu, Fen Wu, Yu Chen, Alejandro Pironti, Huilin Li

**Affiliations:** Division of Biostatistics, Department of Population Health, New York University School of Medicine, New York, New York 10016, USA; Department of Microbiology, New York University School of Medicine, New York, New York 10016, USA; Department of Biological Sciences, Columbia University in the City of New York, New York, New York 10027, USA; Division of Epidemiology, Department of Population Health, New York University School of Medicine, New York, New York 10016, USA

**Keywords:** Microbiome, longitudinal metagenomic data, strain-level analysis, genomic variants, strain dynamics

## Abstract

The development of sequencing technology and analytic tools have advanced our insights into the complexity of microbiome. Since different strains within species may display great phenotypic variability, studying within-species variations enhances the understanding of microbial biological processes. However, most existing methods for strain-level analysis do not allow for the simultaneous interrogation of strain proportions and genome-wide variants in longitudinal metagenomic samples. In this study, we introduce LongStrain, an integrated pipeline for the analysis of metagenomic data from individuals with longitudinal or repeated samples. Our algorithm improves the efficiency and accuracy of strain identification by jointly modeling the strain proportion and genomic variants in combined multiple samples within individuals. With simulation studies of a microbial community and single species, we show that LongStrain is superior to three extensively used methods in variant calling and proportion estimation. Furthermore, we illustrate the potential applications of LongStrain in the real data analysis of The Environmental Determinants of Diabetes in the Young (TEDDY) study and a gastric intestinal metaplasia microbiome study. We investigate the association between the dynamic change of strain proportions and early life events, such as birth delivery mode, antibiotic treatment, and weaning. By joint analysis of phylogeny and strain transition, we also identify a subspecies clade of *Bifidobacterium longum* which is significantly correlated with breastfeeding.

## Introduction

With the steady growth of longitudinal microbiome studies, microbiome is now on the cusp of clinical utility for several diseases, including obesity (John and Mullin 2016; Maruvada et al. 2017), diabetes (Tilg and Moschen 2014; Vallianou et al. 2018), inflammatory bowel disease (IBD) (Manichanh et al. 2012; Ni et al. 2017), and cancer (Karpiński 2019; Vivarelli et al. 2019). The characterization of microbiome was traditionally limited to taxonomic classifications of 16S rRNA sequences (Caporaso et al. 2010). However, the advances in metagenomic sequencing technology and analytic tools have facilitated precise interpretations of the microbial profile and function at species or strain level (Garud and Pollard 2020). Although species level usually is the taxonomic category with the sufficient resolution in the output of metagenomics sequencing data processing pipeline, different strains within species may contain extreme phenotypic variability (Van Rossum et al. 2020). Until now, strain-level analysis has uncovered a broad strain-level diversity within a single person (Zhao et al. 2019), and important implications for antibiotic resistance (Mazel et al. 2000) and pathogen virulence (Leatham et al. 2009). Moreover, multiple coexisting lineages and signal of within-patient selection were observed for *Burkholderia dolosa* (Lieberman et al. 2014) and *Staphylococcus epidermidis* (Lee et al. 2018). Hence, investigating within-species variation holds great scientific promise in understanding the molecular mechanism of microbial biological processes. Extensive efforts have been made to develop tools for analyzing the diversity within species using metagenomic data. PhyloPhlAn 3 focuses on the phylogenetic analysis of multiple strains using clade specific maximally informative markers (Asnicar et al. 2020). StrainPhlAn3 identifies the dominant strains of target species (usually a strain is deemed dominant when its relative abundance exceeds 80%, see Results section for more discussion) and constructs a phylogenetic tree based on predefined MetaPhlAn markers (Truong et al. 2017). PanPhlAn classifies strains based on the entire gene set of the species’ pangenome and characterizes their functional potential in combination with metatranscriptomics (Scholz et al. 2016). Strain Finder (Smillie et al. 2018) infers strain abundances and genotypes using an expectation-maximization algorithm based on 31 single-copy phylogenetic markers from the AMPHORA database (Wu and Eisen 2008). ConStrains discriminates different strains by using barcode-like string of concatenated single nucleotide polymorphisms (SNPs) spanning hundreds of genes (referred to as the ‘uniGcode’) and constructs candidate models of strain combinations using a clustering algorithm (Luo et al. 2015). A common feature of the above methods is their focus on core genes or genetic markers, which may disregard genes that are discriminative of strains, e.g. accessory virulence and drug resistance genes. MIDAS is an integrated pipeline for quantifying microbial species abundance and intra-species genomic variation from shotgun metagenomes in a genome-wide scale (Nayfach et al. 2016). Although MIDAS is an efficient tool for quantification of strain-level gene content and single nucleotide variants (SNVs), it requires >10× and >5× sequencing depth for identification of SNVs and copy number variants in abundant species respectively. There are also many other tools developed for various study designs and clinical applications, which have been well reviewed (Anyansi et al. 2020). However, most methods do not allow for the simultaneous interrogation of strain proportions and genome-wide variants in massive longitudinal metagenomic samples.

The exactly same strains of a species are frequently shared across single-patient microbiome samples that are obtained from different body sites or longitudinally (Yassour et al. 2018; Zhao et al. 2019). In such cases, we may not only be interested in the genomic variants of dominant strains, but also in the genomic variants of non-dominant strains and the dynamic change of these strains’ proportions. Thus, it is critical to jointly model the proportions and genetic variants of strains in longitudinal samples by utilizing raw sequencing data. There are multiple merits in doing this. Firstly, this allows us to more accurately detect the variants of the dominant strain and non-dominant strain (if present). Some methods mainly focus on a dominant strain or a major allele (Truong et al. 2017). For example, Garud’s pipeline (Garud et al. 2019) built on MIDAS required quasi-phaseable (QP) samples with major allele frequencies >80% in the core genome, which neglects a large number of non-QP samples. When two strains have comparable proportions, it is challenging to assign variants correctly to the strains. However, if multiple samples from one subject are available, allele information from other samples can be leveraged to differentiate the genomic variants between the dominant and the non-dominant strains. Secondly, variants that are assigned correctly to each strain will benefit the estimation of strain proportion in return, as genome-wide variants can greatly improve the accuracy of strain proportion compared to the one estimated based on marker genes. Thirdly, we can monitor the dynamic change of strains by modeling strain transitions within species using longitudinal samples (Vatanen et al. 2018; Yassour et al. 2018; Vatanen et al. 2019).

In this paper, we developed LongStrain, a strain-level microbial community profiling pipeline utilizing longitudinal metagenomic data for the detection of genome-wide variants, as well as strain identification and proportion estimation. LongStrain is robust and efficient in performing strain-level analysis within various species from complex microbial communities. The output of LongStrain consists of the genome-wide variants in variant call format and proportion estimates for two strains at all time points. Thus, we can characterize dynamic changes of different strains of microbial species and compare their genomic variations not only in longitudinal samples but also across multiple studies. To validate the performance of our approach, we designed extensive simulations to compare the results of LongStrain with three broadly used tools: StrainPhlAn3, MIDAS, and ConStrains. Additionally, we tested LongStrain on a subset of The Environmental Determinants of Diabetes in the Young (TEDDY) study (Vatanen et al. 2018) and on a study of oral and gastric microbiome in relation to gastric intestinal metaplasia (Wu et al. 2021).

## Results

### An integrated strain-level analytic pipeline using longitudinal metagenomic data

LongStrain is an integrated pipeline for quantifying genomic variants at strain level using metagenomic data from individuals with longitudinal or concurrent samples (Fig. 1). For each subject, raw reads from multiple samples are first assigned to each species by Kraken2 (Wood et al. 2019), which is an efficient tool for taxonomic classification of metagenomic data with both high precision and sensitivity (Simon et al. 2019; Lu and Salzberg 2020). Since reads from one strain may be assigned to several closely related strains, reads that were assigned sub-species designations are aggregated to the species level according to their taxonomy IDs. For each species, the aggregated reads are mapped to the *reference genome* from the NCBI Reference Sequence (RefSeq) database by Bowtie2 (Langmead and Salzberg 2012). If no *reference genome* exists, a *representative genome* or other complete genome (which can be provided by users) will be used. Consequently, our method requires a complete genome of the target species. The input files of our variant calling algorithm are sequence alignment files generated by Bowtie2.

**Fig 1.**
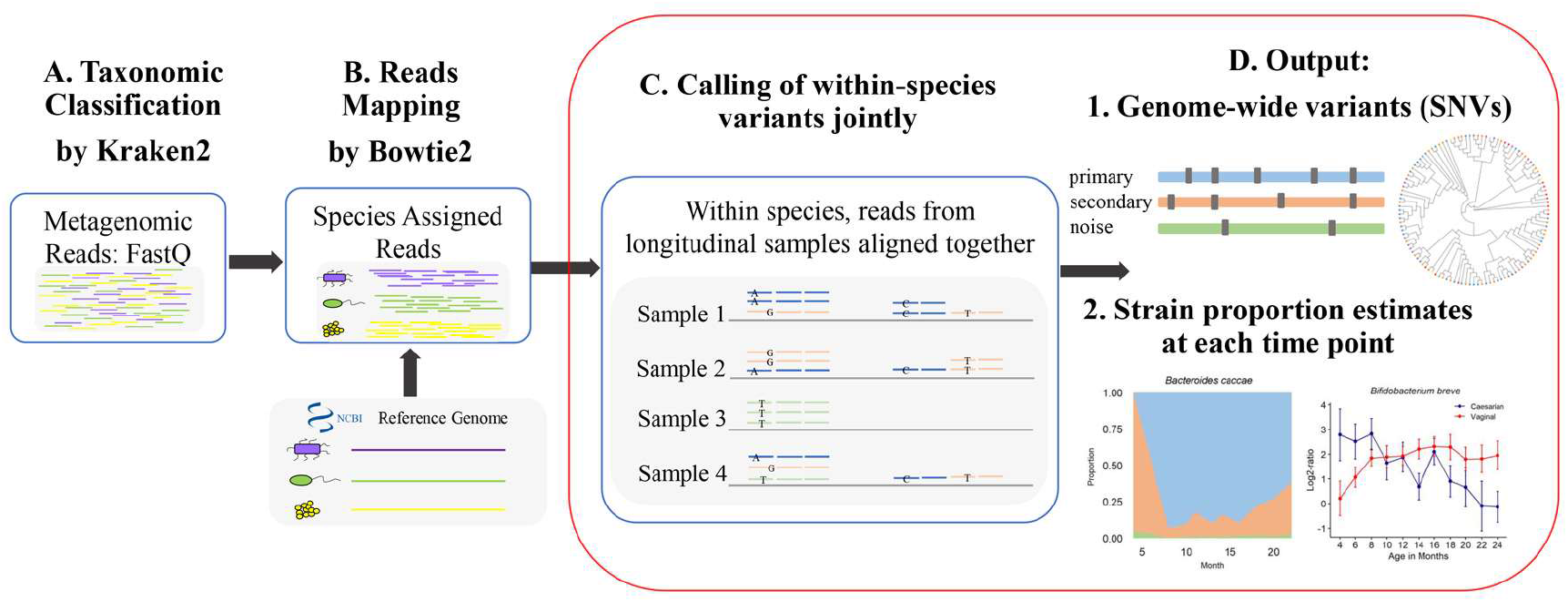
The LongStrain pipeline. (A) Raw reads are assigned to each species by Kraken2. (B) Assigned reads are aligned to the complete genomes extracted from NCBI RefSeq or provided by users. (C) Within each species, genomic variants and strain proportions are inferred jointly by our algorithm in longitudinal or concurrent samples. (D) LongStrain outputs genomic variants and proportions of the primary and secondary strain in longitudinal or concurrent samples.

LongStrain relies on two assumptions. First, for most species, there is one dominant strain within subjects. Second, the vast majority of SNVs in a particular strain are stable within an individual in a short period of time. The first assumption is supported by recent human gut microbiome studies (Truong et al. 2017; Garud et al. 2019). Due to the low mutation rate of DNA-based genomes (Duffy et al. 2008), the second assumption is reasonable in a relative short period of time if horizontal gene transfer or recombination can be disregarded. For example, a culture-based study spanning 2 years estimated the mutation rate of *Bacteroides fragilis* as about 0.9 SNPs/genome/year (Zhao et al. 2019). Even for hypermutating sublineages, the number of novel mutations can be neglected when compared to the total number of variants (Zhao et al. 2019).

In a longitudinal study, the dominant strain at one time point may not be the dominant one at another time point. In order to model the transition of the dominant strains at different time points, we consider three strains in our model, namely a primary strain, a secondary strain, and a noise strain. Here, we define the primary (secondary) strain as the strain with the highest (second highest) proportion of reads within the target species after combining all longitudinal samples from a subject. The genomic variants of primary and secondary strains are assumed to be consistent across multiple samples from one subject and variants that cannot be classified are regarded as being from the noise strain.

For each given species in each subject, LongStrain screens the whole genome of the species to identify strains’ genomic variants and construct their haplotypes. After obtaining sequence alignment files from multiple time points, we pile up all reads from these files together using pysam (Li et al. 2009). Then, the piled reads and variants are classified to the corresponding strains according to our maximum likelihood model (Methods). Note that our model does not account for time information and treats longitudinal samples in the same way as concurrent samples. By pooling microbial genomic data from the same subject together, we increase our power to identify genetic variants and estimate strain proportions.

### Performance assessment using simulated microbial communities (Gut20)

To evaluate the performance of LongStrain, we compared the results of LongStrain with three extensively used reference-based tools: StrainPhlAn, MIDAS, and ConStrains. We simulated a human gut microbial community consisting of 20 species mostly from the Gut20 scenario in Ounit’s study (Kuleshov et al. 2016; Ounit and Lonardi 2016). For each species, we simulated raw sequencing reads (at depth of 10×) from two reference genomes to mimic primary and secondary strains and designed multiple scenarios of longitudinal fluctuations of strain compositions at three time points (Methods). At each time point, the simulated data of all 20 species were pooled to obtain the simulated samples of a microbial community. Finally, all four methods were applied to the simulated dataset. We assessed the performance of SNVs identification using precision and recall (Methods) but omitted ConStrains from this comparison because it uses the marker gene database in MetaPhlAn2 (which has been improved and upgraded greatly in StrainPhlAn3) and it cannot flexibly use other genome databases as references. For the estimation of strain proportions, LongStrain was only compared with ConStrains because the other two tools do not provide this result.

We assessed the performance of the methods using 20 repetitions of the simulation procedure. Concerning the comparison of SNV calling, we tabulated the number of species in which the SNV-calling precision of the primary strain was larger than 0.9 along with their corresponding recall (Methods). On average, LongStrain reported 16 out of the 20 species in each repetition, while MIDAS and StrainPhlAn only reported 10 and 8 out of the 20 species respectively (Fig. 2A). The distribution of average recall of primary strains in all repeats is shown in Fig. 2B, which indicates that LongStrain outputs primary strains in most species with highest recall. With its limited marker-based database, StrainPhlAn unsurprisingly had the lowest recall.

**Fig 2.**
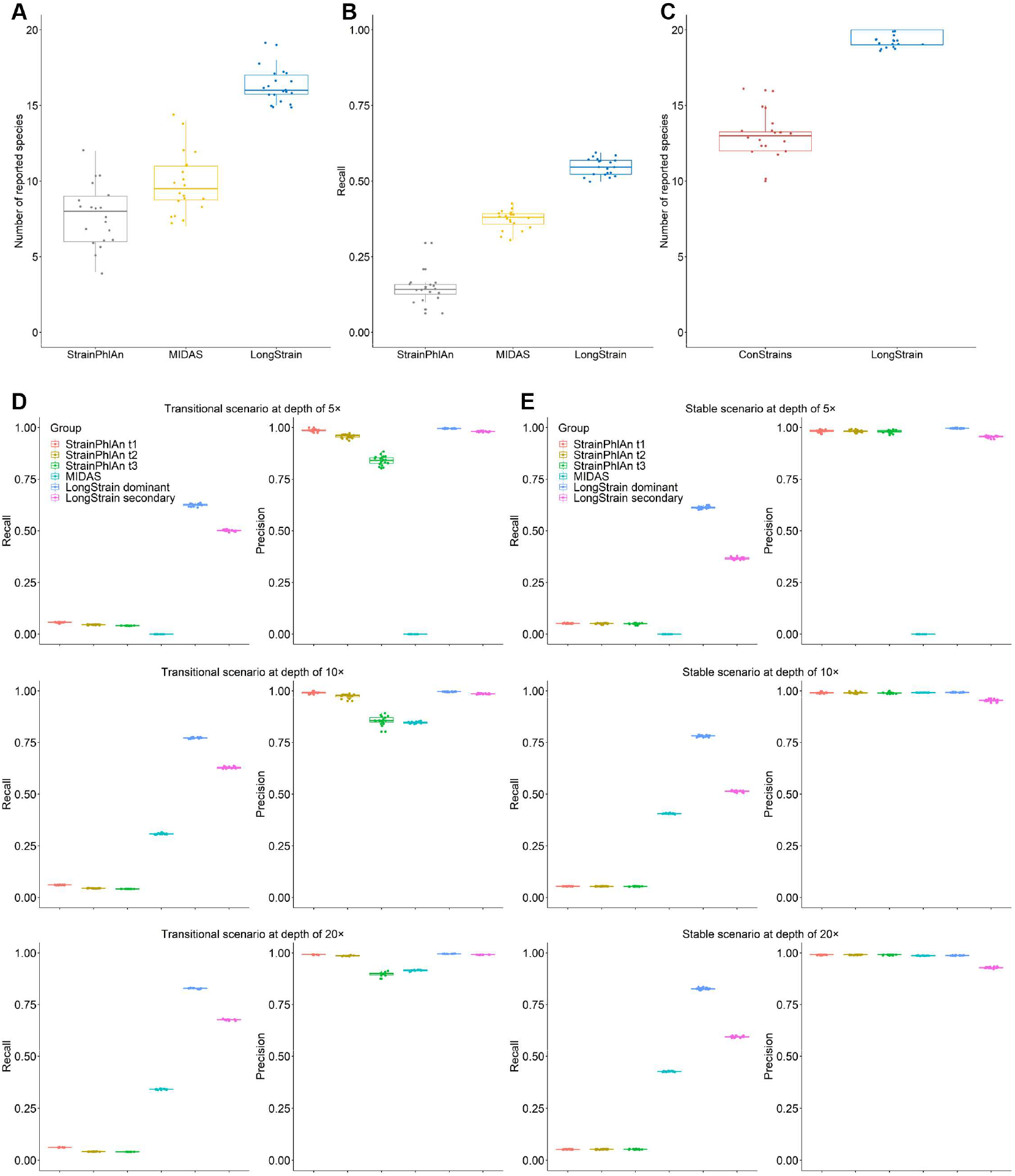
Comparison of LongStrain, StrainPhlAn, MIDAS, and ConStrains in community simulation (Gut20) by 20 repetitions. (A) Number of species in which the SNV-calling precision of the primary strain is >0.9 reported by three methods. (B) Corresponding SNV-calling recall, the proportion of true variants that are correctly identified (in species with precision >0.9) by three methods. (C) Distribution of the number of species successfully reported by two methods respectively in each repetition. (D) Recall and precision of StrainPhlAn, MIDAS, and LongStrain in *Bifidobacterium breve* at average depth of 5×, 10×, and 20× under a transitional scenario. (E) Recall and precision of StrainPhlAn, MIDAS, and LongStrain in *Bifidobacterium breve* at average depth of 5×, 10×, and 20× under a stable scenario.

For the estimation of strain proportions, we compared the primary-strain proportions estimated by LongStrain and ConStrains at three time points. Since there were only two strains simulated in the data, the result was regarded as an unsuccessful estimation if a third strain was estimated to have a proportion larger than 2%. On the simulated dataset and averaged across 20 repetitions, ConStrains only reported the proportions of strains in 13 out of the 20 species, while LongStrain reported more than 19 species (Fig. 2C). We measured the accuracy of the estimated proportions by mean absolute error (MAE) and root mean square error (RMSE) at three time points in each species (Methods; Table 1). LongStrain had higher accuracy in 15 out of 18 species that were successfully reported by ConStrains. In summary, our simulation results show that LongStrain has a much higher overall performance than ConStrains in estimating strain proportions.

**Table 1.** The accuracy of the estimated proportion of the primary strain by ConStrains and LongStrain.

| Species | ConStrains |  |  | LongStrain |  |  |
| --- | --- | --- | --- | --- | --- | --- |
|  | N* | MAE | RMSE | N | MAE | RMSE |
| *A. baumannii* | 8 | **0.048** | **0.060** | **20** | 0.082 | 0.088 |
| *B. cereus* | 0 | - | - | **20** | **0.091** | **0.130** |
| *B. fragilis* | 18 | 0.055 | 0.073 | **20** | **0.028** | **0.039** |
| *B. adolescentis* | 18 | 0.053 | 0.064 | **20** | **0.015** | **0.022** |
| *B. breve* | 7 | 0.047 | 0.052 | **20** | **0.014** | **0.021** |
| *B. longum* | 15 | 0.073 | 0.088 | **20** | **0.012** | **0.017** |
| *C. beijerinckii* | 5 | 0.342 | 0.342 | **20** | **0.045** | **0.049** |
| *D. radiodurans* | 15 | 0.180 | 0.205 | **20** | **0.096** | **0.102** |
| *E. faecalis* | 14 | 0.064 | 0.073 | **20** | **0.019** | **0.026** |
| *E. coli* | 19 | **0.039** | **0.044** | **20** | 0.219 | 0.232 |
| *H. pylori* | 18 | 0.098 | 0.104 | **20** | **0.041** | **0.048** |
| *L. gasseri* | 0 | - | - | **6** | **0.165** | **0.173** |
| *L. monocytogenes* | 12 | 0.067 | 0.071 | **20** | 0.067 | **0.069** |
| *N. meningitidis* | 13 | 0.042 | 0.051 | **20** | **0.035** | **0.038** |
| *P. aeruginosa* | 19 | 0.047 | 0.052 | **20** | **0.031** | **0.038** |
| *R. sphaeroides* | 19 | 0.077 | 0.086 | **20** | **0.021** | **0.022** |
| *S. aureus* | 17 | 0.049 | 0.056 | **20** | **0.024** | **0.028** |
| *S. epidermidis* | 18 | 0.058 | 0.064 | **20** | **0.014** | **0.017** |
| *S. agalactiae* | 19 | **0.063** | **0.068** | **20** | 0.117 | 0.132 |
| *S. mutans* | 9 | 0.034 | 0.038 | **20** | **0.024** | **0.029** |

\*: N is the number of successful reports in 20 repetitions of the simulation. Values in bold highlight the better performing method for a given species.

### Performance assessment using simulated reads from single species

Considering that the inherent characteristics of microbe species, such as prevalence, pathogenicity, genomic complexity, and the longitudinal scenario can affect the performance of microbiome analysis methods, we simulated and analyzed the data of the 20 species used in the above community simulation separately (Methods) in order to understand how the different factors affect LongStrain. For most of these species, we considered two representative temporal scenarios. One is a transitional scenario in which the ratio of the sequencing depths of the primary strain vs. the secondary strain changes from 9×:1× at the first time point to 6×:4× and 3×:7× at two later time points respectively. In the other stable scenario, the ratio of the sequencing depths of two strains remains constant, 8×:2×, across three time points.

In particular, we tested more scenarios with different depths in *Bifidobacterium breve* (Methods), for which all methods had stable performances. We compared the recall and precision of three methods at average depth of 5×, 10×, and 20× in the transitional (Fig. 2D) and stable (Fig. 2E) scenarios. With incremented average depth, all methods had better performances as expected. Since MIDAS requires a minimal depth of ~10× to identify the majority of SNPs, its recall and precision at the depth of 5× were marked as zero. In contrast, LongStrain had a relatively high recall and precision for the primary strain and the secondary strain when the depth was as low as <5× at a single time point. We also confirmed the similar effect of depth in *B. fragilis* (Supplemental Fig. S1).

Since MIDAS and StrainPhlAn only take into account the primary strain, they display a higher precision under the stable scenario compared to the transitional scenario (Fig. 2D, E). Especially at high sequencing depths, they show a slightly higher precision on the primary strain than LongStrain (Supplemental Figs. S2, S3 show performance assessment with *Bifidobacterium adolescentis* and *Staphylococcus aureus*). In contrast, LongStrain makes a trade-off between the accuracy of the primary strain and the secondary strain. For most species, LongStrain displays a lower recall and precision on the second strain than on the primary strain due to the former’s lower proportion of reads. In *Enterococcus faecalis* and *Listeria monocytogenes*, however, the recall of the secondary strain is higher than that of the primary strain (Supplemental Figs. S4, S5), which may be caused by the higher genetic distance of the secondary strain to the reference genome than that of the primary strain (Supplementary Table 1).

We also compared the proportions of strains estimated by LongStrain and ConStrains in each species under two scenarios. Similar to the results of the community simulation, the proportions of reported species by ConStrains in each repeat are around 65% and 90% respectively under two scenarios (Supplemental Fig. S6). The reporting rates of LongStrain are consistently close to 100%. In terms of the bias of the estimated proportions, LongStrain still performed better in more species than ConStrains, though ConStrains had smaller bias in some species (Supplemental Tables S2, S3).

In summary, LongStrain outperformed the three competing methods in most species with high-quality reference genomes, especially at relatively low sequencing depths (5~10×). LongStrain also performed better than other methods for a potential pathogen, *Helicobacter pylori* (Supplemental Fig. S7), with numerous closely related genomes in the database. However, LongStrain could not get reliable genomic variants in *Escherichia coli* due to the errors introduced in reads classification and mapping, for which reason *E. coli* was omitted in part of the real data analysis related to the variant calling.

### Using LongStrain to obtain further insights from previous microbiome studies

We applied LongStrain on two datasets from previously reported microbiome studies. The first dataset stems from the TEDDY study and we use it to showcase four major applications of LongStrain in metagenomics research. LongStrain can: 1) provide longitudinal proportion estimates of strains within species, which enable temporal pattern analysis at strain level; 2) identify strain transitions, which allows strain transition time to be compared to other time-dependent factors; 3) study diversity at the strain level; and 4) conduct phylogenetic tree-based strain-level association tests, which enable the identification of strains or species subclades related to the relevant clinical outcome. The second real data come from a study of the oral and gastric microbiome in relation to gastric intestinal metaplasia, with which we add another interesting example of phylogenetic tree-based strain-level association tests (the fourth application in the above list).

### Real data 1: TEDDY metagenomics study

We first retrieved a set of 100 subjects with T1D diagnosis and 100 controls from the TEDDY project (Vatanen et al. 2018) including six clinical research centers in the United States (Colorado, Georgia/Florida and Washington) and Europe (Finland, Germany and Sweden). We applied LongStrain to the dataset and screened 44 species (Supplemental File) with relatively high abundance reported in two previous TEDDY studies (Stewart et al. 2018; Vatanen et al. 2018). After filtering, there were 26 species which had sufficient sequencing depth (sum of depth >10× in the longitudinal samples) retained for the strain-level analysis.

### Application 1: Longitudinal analysis of strain proportion

We denote the proportion of a strain within its species at one time point as *p*. The dominant strain at a certain time point is defined as the strain with p>80% (Garud et al. 2019). Note that, the “dominant” strain is defined at each time point (or for each sample), which is different from the “primary” strain previously defined in the combined longitudinal samples. Most samples have a dominant strain (Supplemental Fig. S8), which agrees with previous studies (Truong et al. 2017; Garud et al. 2019). However, there is still a considerable proportion (ranging from 15.2% to 67.4%) of samples in which the highest *p* is <80% in each of these 26 species, which emphasizes the necessity to take non-dominant strains into account. To illustrate the potential applications of estimating strain proportions, we calculated the log2-ratio of the proportions of the primary strain to the secondary strain. Then, we tested whether the temporal trend of the log2-ratio was associated with the birth delivery mode (caesarian or vaginal) using a linear mixed model (LMM) adjusting for gender (Methods). In Supplemental Table S4, we identified five species in which the temporal trends of the log2-ratio were significantly different between two birth delivery modes after false discovery rate (FDR) correction: *B. breve* (q=5.79E-08), *Akkermansia muciniphila* (q=0.020), *Bacteroides thetaiotaomicron* (q=0.020), *Faecalibacterium prausnitzii* (q=0.034), and *Roseburia intestinalis* (q=0.034). Shown as an example in Fig. 3A, the log2-ratio increases in infants born by vaginal delivery but decreases in infants born by cesarean in *B. breve*.

**Fig 3.**
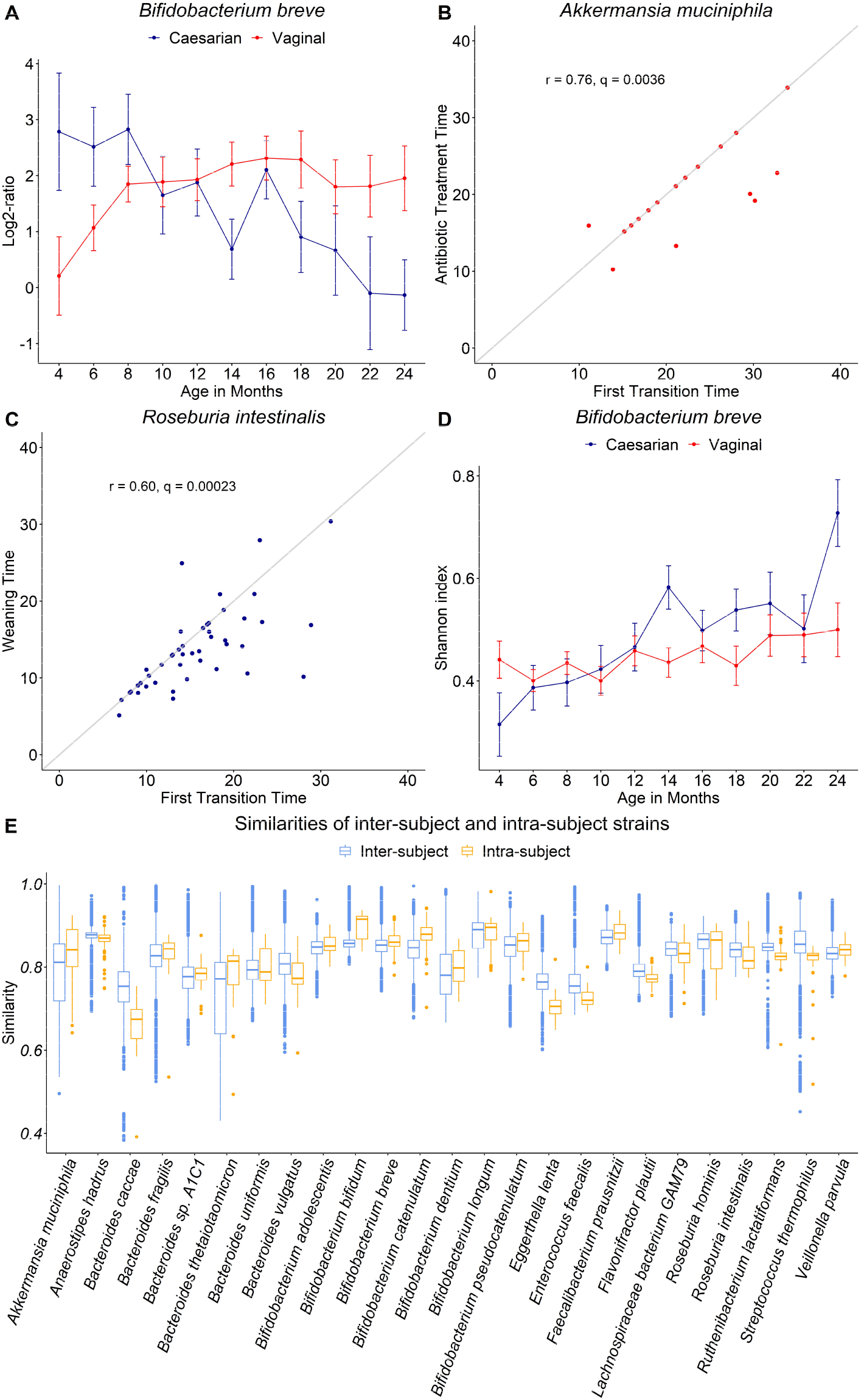
A. Log2-ratio of the proportions of the primary strain to the secondary strain by birth delivery mode in *B. breve*. B. Scatter plot and Pearson correlation test of time (in months) of the first transition and the antibiotic treatment in *A. muciniphila*. C. Scatter plot and Pearson correlation test of time (in months) of the first transition and the weaning in *R. intestinalis*. D. Shannon index of the strains by different delivery modes within *B. breve*. E. Variants similarity of inter-subject and intra-subject strains in 25 species.

### Application 2: Identifying strain transitions

To explore the transition of dominant strains across time, we define the major or minor strain as the strain with p>=50% or p<50% in one sample, respectively. If the major strain becomes minor in the longitudinal samples of a subject, we call it a strain transition, which may indicate a drastic change of microenvironment. With the above definitions, we investigated whether a transition happened in subjects for the species mentioned above at time points with sufficient read counts. We found that strain transition was prevalent in most species in the early gut microbiome (months 3 to 46) (Supplemental Fig. S9). The proportion of subjects with a strain transition ranges from 18.4% (in *Bacteroides thetaiotaomicron*) to 96% (in *Lachnospiraceae bacterium GAM79*) in 26 species.

Next, we focused on the subjects who had early life events (antibiotic treatments and weaning) and at least one strain transition to explore their relationship. We performed the Pearson correlation test and drew the scatter plot of the times of the first transition and early life event. After FDR correction, we observed significant correlations between the time of the first transition and the time of receiving antibiotics in *A. muciniphila (r=0.76*, q=0.0036), *Bifidobacterium bifidum* (r=0.69, q=0.0036), *Bifidobacterium longum* (r=0.40, q=0.0072), and *E. coli* (r=0.48, q=0.0036) (Fig. 3B, Supplemental Fig. S10 A, B, C). Particularly in *A. muciniphila*, the first strain transitions coincided with the antibiotic treatments in subjects, which was indicated by the points on the diagonal line (Fig3. B). Weaning was found to be correlated with the first transition in *R. intestinalis* (r=0.60, q=0.00023), *B. bifidum* (r=0.54, q=0.0021), *B. longum* (r=0.42, q=0.00086), and *Roseburia hominis* (r=0.71, q=0.0031) (Fig 3. C, Supplemental Fig S10. D, E, F).

### Application 3: Analysis of intra- or inter-subject diversity at strain level

With the proportions of strains outputted from LongStrain, we calculated the Shannon’s index of strain proportions within species to measure the strain diversity (Method). Similar to the log2-ratio of strain proportions, we also tested whether the temporal trend of Shannon’s diversity over time was associated with two birth delivery modes. A significant association was only observed in *B. breve* (q=0.0072) by the analysis of LMM adjusting for gender (Method). Vaginally delivered infants has a higher temporal variation of strain-level diversity in *B. breve* than cesarean-born infants (Fig. 3D), which is consistent with the delayed colonization of *B. breve* in C-section-delivered infants (Chua et al. 2017).

With the genomic variants outputted from LongStrain, we then extracted the polymorphic sites across strains within each species since we obtained the genomic variants of primary and secondary strains. The variant similarity between two strains was defined as the percentage of identical genotypes at polymorphic sites. We stratified the similarities by intra- and inter-subject comparison in 25 species (*E. coli* was omitted, Fig. 3E). The former was the similarity between the primary and the secondary strain within each subject and the latter was the similarity between strains across different subjects. The median similarity in most species ranged from 70% to 90%, and the median intra-subject similarity is close to that of inter-subject similarity in most species, which indicates that two strains within a subject do not tend to have a higher variant similarity than strains from different subjects.

We were interested in investigating whether the similarity of variants was related to the case-control status. Specifically, if the strains within cases or within controls are genetically more uniform, this should manifest as a higher variant similarity within either of the groups. To test this hypothesis, we assigned the similarities between all pairs of strains to three groups, “*case vs case*”, “*control vs control*”, and “*case vs control*”. Then, we obtained the distributions of similarity values within each group. To eliminate the effect of the primary or secondary strain, the above analysis was repeated within pairs of primary strains and pairs of secondary strains respectively. However, we did not find significantly different distributions of similarity between “*case vs case*” and “*control vs control*” groups after screening 26 species above. The conclusion did not change when we focused on the analysis of primary strains and secondary strains separately. For example, the results of *A. muciniphila*, *B. uniformis*, and *B. longum* are shown in Supplemental Figs. S11, S12, S13. It may imply that the onset of T1D is not associated with phylogenetic clusters of strains for these species.

### Application 4: LongStrain in phylogenetic tree-based strain-level association test

We further explored the significant factors associated with phylogenetic distances between identified strains. We constructed neighbor-joining trees of species with MEGA-X (Kumar et al. 2018) using variants identified by LongStrain. Since the TEDDY cohort was largely non-Hispanic white, it was not surprising that strains did not cluster by centers on the phylogenetic trees. Specifically, most species did not show discrete population structure within Westernized populations (for example, Supplemental Fig. 14), in accordance with a previous study (Truong et al. 2017). Nonetheless, we observed some clusters of strains by geographic location in *B. breve* and *B. longum*. Particularly in *B. breve* (Fig. 4A), strains highlighted in pink were almost exclusively from three centers (Colorado, Georgia, and Washington) in the USA and subclades highlighted in blue mostly consisted of strains from two Northern European countries (Finland and Sweden). Conversely, subclades in green showed diversified sources of strains. Additionally, similar patterns were observed in the phylogenetic tree of *B. longum* (Fig. 4B). Although not as distinct as those in *B. breve*, some strains from the USA or Northern European countries clustered under some subclades of *B. longum*. To further explore whether subclades were correlated with other factors, we tested the association between the phylogenetic patterns of strains and T1D diagnosis, delivery mode, antibiotics, breastfeeding, and phases of microbiome progression separately. Intriguingly, we observed that only a subspecies clade of *B. longum* was associated with breastfeeding or phases of microbiome progression (Fig. 4C). The status (primary or secondary) of each strain was recalculated before and after breastfeeding. The majority of the subspecies clade highlighted in pink (Fig. 4 B and C) displayed a switch from “primary” to “secondary”. The microbiome progression in early life was previously defined as three phases, a developmental phase (months 3–14), a transitional phase (months 15–30), and a stable phase (≥31 months) according to the Shannon’s diversity index (Stewart et al. 2018). The status (primary or secondary) of each strain was recalculated in each phase. Naturally, a similar transition of strain status was observed from the first phase to the third phase for this clade (Fig. 4 D). This phylogenetic pattern coincided with the well-characterized subspecies clade of *B. longum*, *B. infantis*, identified in the DIABIMMUNE study in Finland, Estonia and Russian Karelia (Vatanen et al. 2019). *B. infantis* is capable of efficiently consuming several small mass human milk oligosaccharides HMOs and harbors a wide variety of genes dedicated to HMO metabolism (Sela et al. 2008; Sela and Mills 2010), which explains its dominance during breastfeeding (or the first phase) and dissipation over time. This analysis highlights that the use of LongStrain empowers us to identify subclades with distinct functionalities in longitudinal metagenomics data.

**Figure 4.**
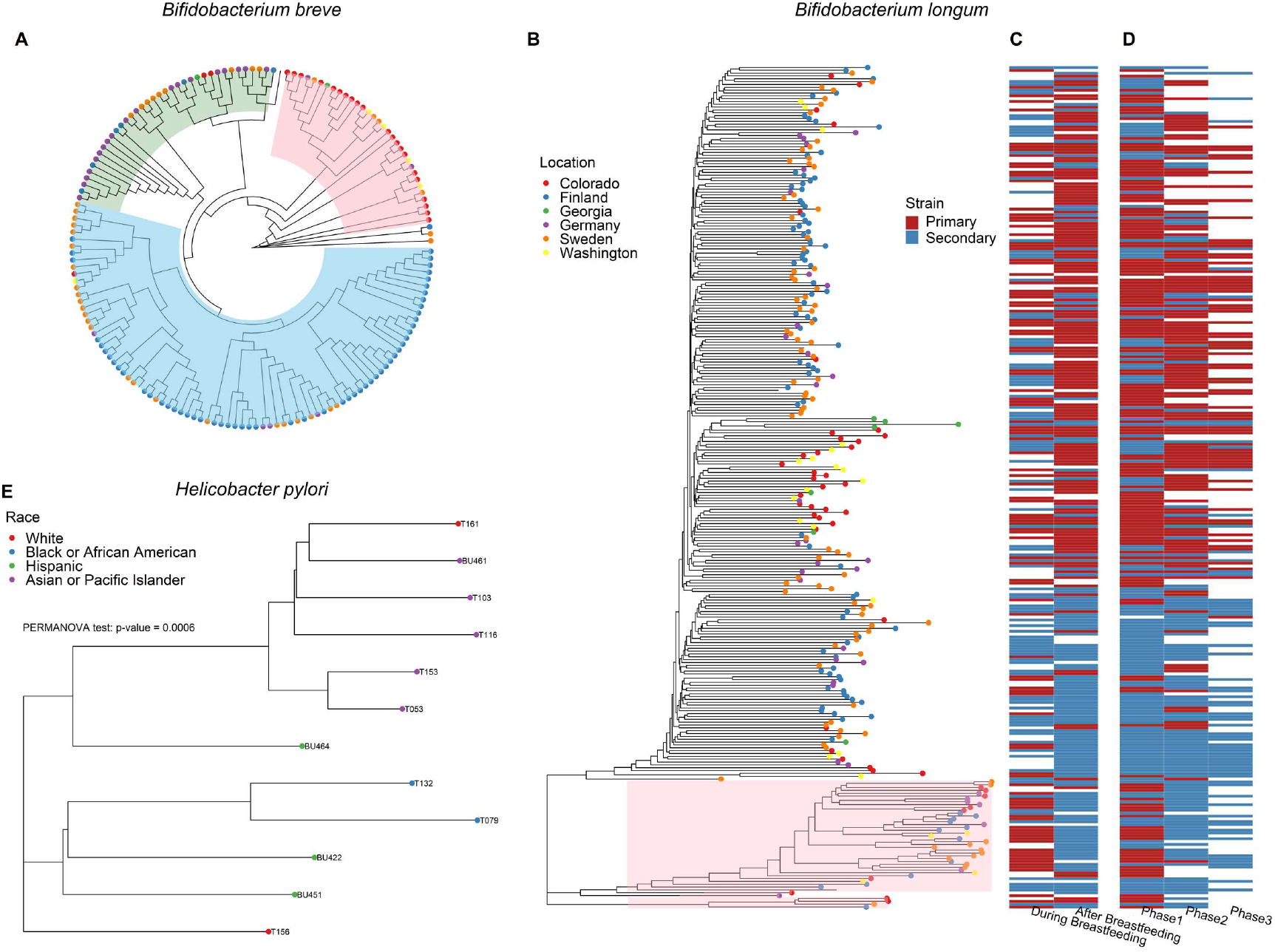
(A) Phylogenetic tree of strains in *B. breve* from six centers. Highlighted in pink: mostly from three centers in the USA; highlighted in blue: mostly from two Northern European countries (Finland and Sweden); highlighted in green: diversified sources from Europe. (B) The phylogenetic tree of *B. longum* in samples with T1D diagnosis from the TEDDY study. The sources of strains are marked by colored dots. The subclades highlighted in pink are associated with breastfeeding and phases of microbiome progression. (C) The status (primary or secondary) of strain before and after breastfeeding. (D) The status (primary or secondary) of strain in phase1 (months 3–14), phase2 (months 15–30), phase3 (≥31 months). (E) Phylogenetic tree of primary strains in *Helicobacter pylori* from 12 subjects.

### Real data 2: a gastric intestinal metaplasia microbiome study (example for Application 4)

We retrieved oral and gastric microbiome data from a case-control study including 89 cases with gastric intestinal metaplasia (IM) and 55 matched controls in New York City, USA (Wu et al. 2021). Oral wash and antral mucosal brushing samples from the same subject were treated as repeated measures in this analysis.

Although *H. pylori*-induced gastritis plays an essential role in gastric cancer initiation, only about 3% infected individuals develop malignancy (Ernst et al. 2006; Polk and Peek 2010). Therefore, in this analysis, we focused on *H. pylori* and sought to identify strains or subclades of *H. pylori* closely related to gastric IM. Unfortunately, due to the fading of *H. pylori* under achlorhydric condition in precancerous lesions (Kwak et al. 2014), *H. pylori* was only detected in 12 subjects with enough sequencing depth (>10×). Since no secondary strain had enough genome coverage (depth <1×), we focused on the primary strains from these subjects in the phylogenetic analysis. Despite the small sample size, the phylogenetic distances between strains were significantly dependent on the race of the hosts (PERMANOVA test, *p*-value < 0.001, Fig. 4 E). We observed the clustering of strains from African Americans and Asians, which was not observed in Hispanics or whites. This may be attributed to the cohesion of community or the dietary habits in these two ethnic groups. On the other hand, the evidence of relatedness between strains and gastric IM was not found in the phylogenetic analysis. For further investigation of the association between gastric cancer and variants in *H. pylori*, more genomes of *H. pylori* with high sequencing depth are needed.

## Discussion

In this study, we introduced an integrated pipeline for both the identification of bacteria genome-wide variants across complex microbial communities and observation of dynamic changes of bacterial abundance at strain level in longitudinal metagenomic data. LongStrain is suitable for metagenomic samples obtained longitudinally or from multiple body sites. By exploiting the genomic variants and proportions of strains in the combined samples from either longitudinal or multiple samples, LongStrain enables accurate characterization of the primary and secondary strains and genetic variation identification. By comparing LongStrain with three popular tools on the simulated datasets, we demonstrated that LongStrain is superior or comparable to these methods in both accuracy and reporting rate under various scenarios. In the real data analysis, we applied LongStrain to a subset of the TEDDY project to extensively characterize prevalent species at strain level. These results are in agreement with previous observations that there is a dominant strain within most species for each sample (Truong et al. 2017). We also observed prevalent transitions of the major strain in some species during the early life of infants, which may be associated with events like antibiotic treatments and dietary changes. We phylogenetically profiled the strains in more than 20 species with the variants identified by LongStrain and found discrete population structures in *B. breve* and *B. longum*. Notably, combining the phylogenetic results with dynamic changes of strains allowed us to directly observe the link between a previously reported subspecies clade of *B. longum* (Vatanen et al. 2019) and breastfeeding.

One of the main advantages of our pipeline is its ability to fully utilize repeated measures and reduce noise by preprocessing. As a result, LongStrain can tackle lower sequencing depth than MIDAS and ConStrains. Owing to the simultaneous estimation of strain proportions and identification of genomic variants, LongStrain can estimate the proportion of the strain of interest directly from repeated samples. In contrast, methods that focus on the dominant strain need to identify genomic variants in different samples separately so that they can determine whether these strains are the same one. Since LongStrain only ensures the accuracy of the primary strain and possible secondary strain, our method is not suitable for samples in which multiple strains have comparable proportions. Processing such samples with LongStrain might yield high estimates (>10%) for the noise strain. Additionally, if the proportions of the primary and secondary strain are consistently close to each other (e.g. 50% to 50%), our method will not correctly assign variants to the strains. Fortunately, this is not a common situation because samples usually have one dominant strain (Truong et al. 2017).

Many tools have been developed for the analysis of metagenomic data at strain level, yet each one of them was designed with a specific scope in minds. Excelling at large-scale strain-level analyses of longitudinal microbiome samples, LongStrain, as a novel method to link strain-level genotypic traits with dynamic changes of strain proportions, addresses an important analysis need. In addition, LongStrain’s ability in accurately identifying strain-level variants offers a solid foundation for further downstream analyses, such as microbial genome-wide association studies and genomic evolution studies.

## Methods

### Preprocessing of repeated metagenomic sequencing samples

As illustrated in Figure 1, in LongStrain, the raw metagenomic sequence reads are first assigned to each species by Kraken2 (Fig. 1A). For each species, based on the selected *reference genome* from the NCBI RefSeq database (Fig. 1B), reads from the same subject are aligned and piled up using pysam (Fig. 1C) (Li et al. 2009). The input to LongStrain consists of species-specific read alignments from the repeated metagenomic sequence samples of one subject. To discriminate between strains, LongStrain focuses on the *effective sites* of the sequence, that is, the genomic sites where the observed major allele frequency in the aligned reads from all the samples is <0.9 or the observed major allele frequency is <0.9 in at least two samples. These filter criteria are designed to reduce noise from sequencing and mapping error.

### Notation preparation

As an example, in Fig. 5, for one individual, there are four longitudinal samples. Within each sample, reads (represented by strips) covering the first three consecutive effective sites 1, 2, and 3 on the genome are aligned. Different colors in strips indicate different strains. According to our assumptions, reads with the same color should have the same genotype at the same effective site (in other words, make up the same haplotypes). As annotated in Fig. 5, we use the letters *i, j*, and *k* to index the effective site, sample, and read respectively. Note that the time order of longitudinal samples is neglected, as LongStrain does not distinguish between longitudinal and parallel samples.

**Figure 5.**
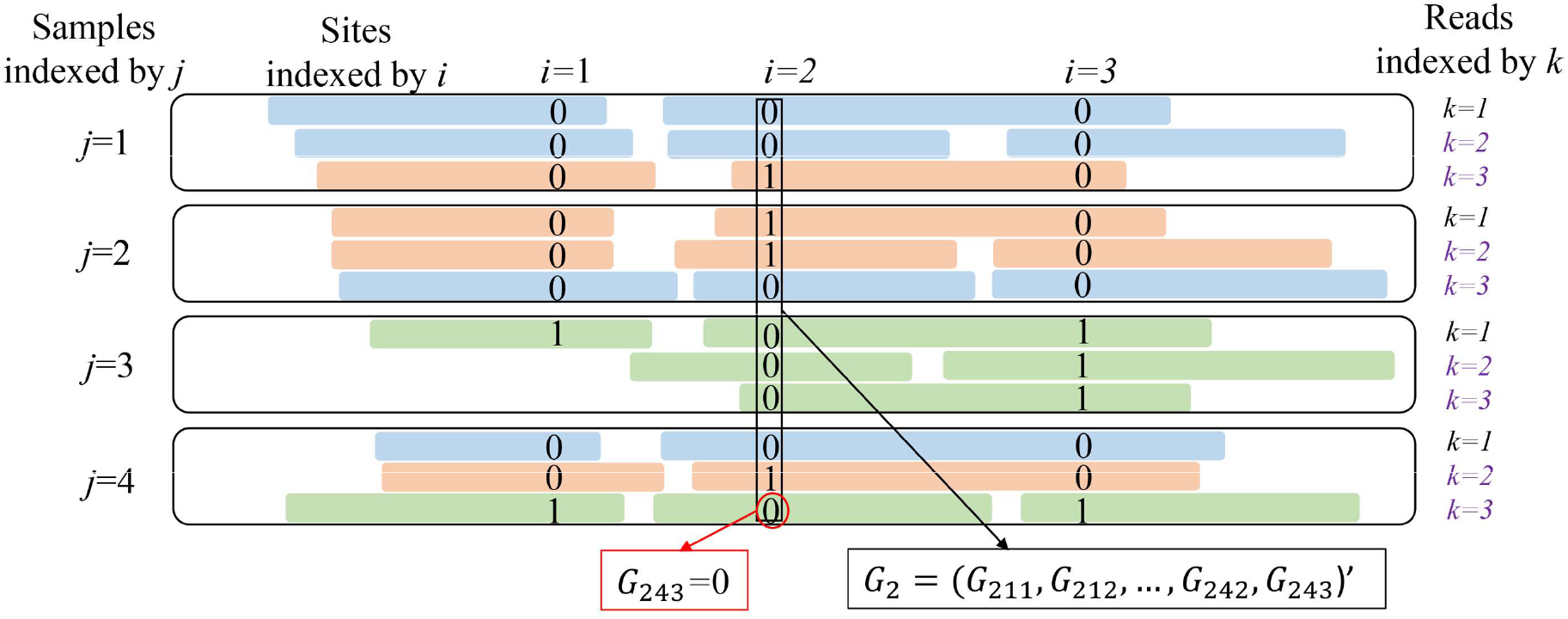
Input data for LongStrain algorithm. For a given species, reads aligned to sites (*i* = 1, 2, 3) from one subject are piled up within four samples (*j* = 1, 2, 3, 4). Strips indicate the reads and different colors indicate different strains. The observed genotype at site *i*, sample *j* and read *k* is denoted by *G_ijk_*.

The first step of LongStrain is to use the observed genotype data to classify the reads to one of three different strain types (primary, secondary or noise). We denote the strain assignment for the *k*^th^ read in the *j*^th^ sample based on the genotype data up to site *i* as *S_ijk_*, and assume that in each sample, there are at most three strains (indexed by *s, s*=1, 2, or 3): primary strain, secondary strain, and noise strain. The noise strain is always assumed to be present in the sample such that the undetermined reads and possible errors are absorbed. The strain assignment is achieved by maximizing the likelihood of the genotypes we observed (denoted as *G_ijk_*) under various possible combinations of strain identification. In the second step, with the reads’ strain assignments, we can estimate the proportion of strain *s* in sample *j* (denoted by *P_js_*) by the ratio of the number of reads classified as strain *s* and the total number of reads in sample *j*, for a certain considered genetic region. We divide the whole genome into windows of 10,000 bp and each window can contain multiple sites. We update the matrix of proportions for the three hypothesized strains in *J* samples, denoted by

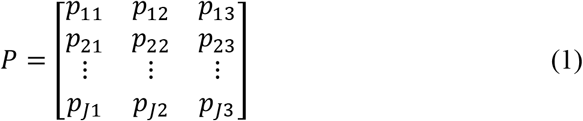

window by window. For the simplicity, we drop the index of windows in all the notations related to *P*. We will introduce how to update *P* in the subsection “**Estimation of strain proportions**”.

### Algorithm of LongStrain

In order to describe the details of LongStrain algorithm, we divide the effective sites into two categories: 1) Non-sharing sites which do not share any reads with other sites in any of samples (e.g. site 2 in Fig. 5). At such sites, only two strains can be differentiated because each site has only two possible genotypes. 2) Sharing sites, which shares some reads with an adjacent site (e.g. site 3 in Fig. 5). At such sites, we are able to differentiate three strains due to the information contained in the reads shared with the adjacent site. Next, we introduce the LongStrain algorithm by site category.

### Category 1: Non-sharing sites

Assuming that there are only two strains present, there are only six possible combinations (indexed by *h* in row) of genotypes (0/1) on the three possible strains (indexed by *s* in column) at such site, denoted by a 6×3 matrix *Θ*,

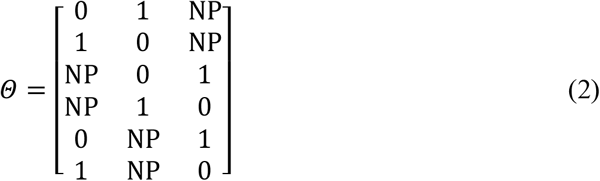

where NP indicates that the strain is not present. For example, given *G_ijk_*, the observed genotype of read *k* from sample *j* at site *i*, by matching it to the *h*^th^ row of *Θ: Θ_h_* = (*Θ*_*h*1_, *Θ*_*h*2_, *θ*_*h*3_)′, we can give the strain identification of this read by:

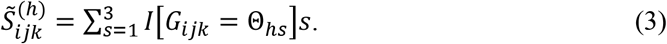

That is, 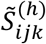 is the strain index of the only strain compatible with the observed genotype. Then we can calculate the likelihood of observing the genotypes of all reads aligned to site *i*, denoted by *G_i_* = (*G*_*i*11_,…, *G*_*i*1*n*_1__,…, *G*_*iJ*1_,…, *G*_*iJn*_J__)′ (as illustrated in Fig. 5) with the *h*^th^ combination as

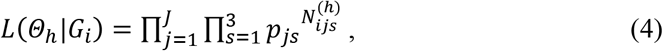

where 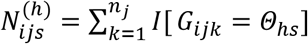 is the number of reads compatible with the given combination of genotypes, *p_js_* comes from the most updated *P* matrix (1), and *n_j_* is the number of reads belonging to sample *j* at each site. In practice, *n_j_* varies across sites, but for simplicity of notation we leave this dependence implicit when describing the method.

We perform the same calculation for *h*=1, 2,…, 6, and find the combination that maximizes the above likelihood (4) and denote this combination with *h*\*. Then the strain assignments of reads covering site *i* is obtained by

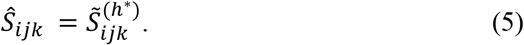

### Category 2: Sharing sites

With sharing of reads between the previous and current sites, we are able to differentiate three strains. Again, there are six possible combinations (indexed by *h* in row) of the genotypes on the three strains (indexed by *s* in column) at such site, denoted by a matrix *Θ*′ with

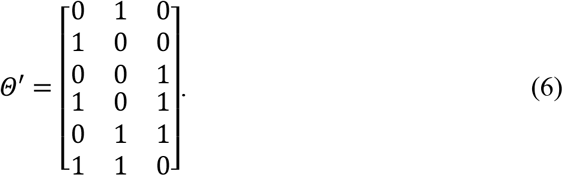

We define 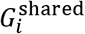 as the genotype of the reads which also cover site *i* – 1 at site *i*, and 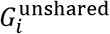 as the genotype of the reads that do not cover site *i* – 1 at site *i*. We use the vector *U_i_* = (*U*_*i*11_,…, *U*_*i*1*n*_1__,…, *U*_*ij*1_,…, *U_ij_j′__*,…, *U_iJ_*,…, *U_iJn_J__*)′ to indicate whether reads covering site *i* also cover site *i* – 1 or not. Specifically, if the read at site *i* also covers site *i* – 1, *U_ijk_* = 1; otherwise *U_ijk_* = 0.

The likelihood of observing *G_i_*, at site *i* given the *h*^th^ combination in *Θ*′ can be calculated as

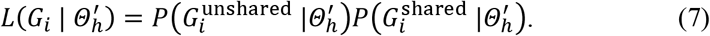

For the unshared reads, the probability of observing their genotypes 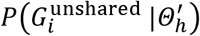 can be calculated similarly to equation (4) in the first category,

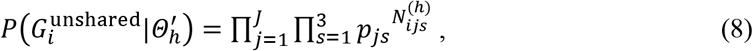

where 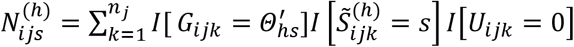 is the number of unshared reads compatible with the given combination of genotypes and strain index *s*, and 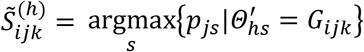. Note that because there are three possible strains (1, 2, or 3) and only two possible genotype values (0/1), there is chance that *G_ijk_* matches to two strains. For example, if *G_ijk_* = 0, for *h*=1, both strain 1 and strain 3 have genotype 0. In such a case, the definition of 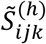 and indicator function 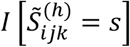 ensure that the read is assigned to the strain whose *p_js_* is larger.

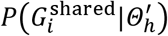 is the probability of observing genotypes of the shared reads. Since the shared reads have already been assigned to a strain at the previous site *i* – 1, we denote the strain identity estimated from the previous site for the *k*^th^ read in the *j*^th^ sample at *i*^th^ site as 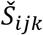. For notational simplicity, when *U_ijk_* = 0, we will arbitrarily set 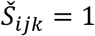 to avoid the undefined value in the below equation (9).

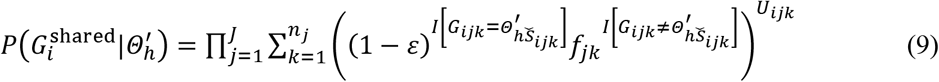

where 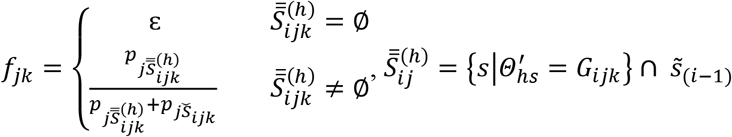

In (9), ε is the error caused by sequencing or mapping and is set to 0.001 (Ma et al. 2019) for simplicity of computation. When 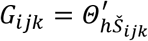, the genotype of read *k* in sample *j* is consistent with the strain identity obtained at the previous site, we keep its strain identity and assign the probability to observe *G_ijk_* as 1 – ε. When 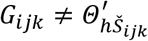, the observed genotype is inconsistent with the inferred strain from the previous site, we need to decide which strain the read belongs and what is the probability to observe the genotype. Let 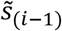 denotes the strain that is not detected at site *i* – 1. For example, if strain-1 and strain-3 are detected at the previous site, then 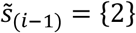. If 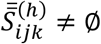 there is one strain that is not detected at the previous site and whose genotype agrees with *G_ijk_*. In other word, read *k* which is assigned to 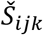 at site *i* – 1 is more likely from 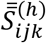. If 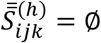, the inconsistency between *G_ijk_* and 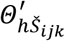 will be attributed to the sequencing or mapping error. We perform the calculation for *h*=1, 2,…, 6 to find the *h* that maximizes the likelihood (7) and denote it as *h*\*. Then the strain identification 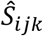 can be obtained according to the following rules,

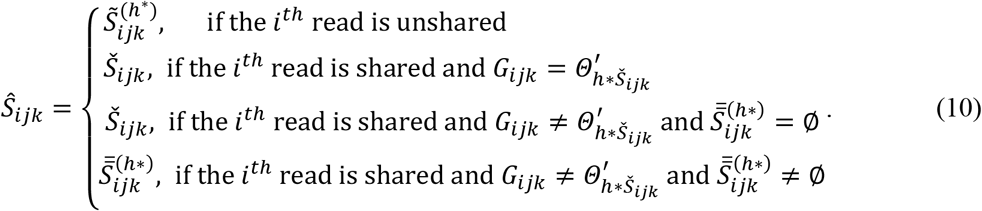

### Estimation of strain proportions

Finally, we introduce how *P* is initialized and updated. We define *C* as the count matrix of reads from three hypothesized strains in *J* samples.

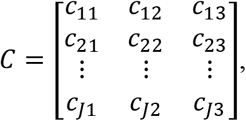

where *C_js_* represents the cumulative count of reads from strain *s* in sample *j. p_js_* is calculated as

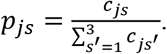

We divided the whole genome into windows of 10,000 bp and calculate the average depth of each window. After sorting all the windows by their average depth, we selected five windows as close as possible to the 80^th^ quantile. We initialized the analysis in these five windows with an equal starting value of 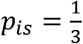. Using the algorithm introduced above, we assigned reads in these regions to three hypothesized strains and obtain the initial *P* and *C*. Then, we screened the whole genome and update *P* and *C* window by window until all are finished.

### Simulation study of microbial community

To compare LongStrain with three competing methods in a more realistic situation, we simulated a gut microbial community (Gut20) which consisted of 20 species used in the previous studies (Kuleshov et al. 2016; Ounit and Lonardi 2016). Since *Actinomyces odontolyticus, Bacteroides vulgatus, Propionibacterium acnes*, and *Propionibacterium acnes* did not have complete genomes in the NCBI database, we replaced them with four other common species in human gut (*B. breve*, *B. adolescentis*, *B. fragilis*, *B. longum*). For each species, we chose two complete genomes (not *reference genome*) at random from the NCBI RefSeq database (for species name and strain, see Supplemental Table S1). To mimic longitudinal changes in the abundance of two strains, we conceived the following scenarios for sequencing depths of two strains at three time points: [9:1, 9:1, 9:1], [8:2, 8:2, 8:2], [7:3, 7:3, 7:3], [6:4, 6:4, 6:4], [9:1, 8:2, 7:3], [9:1, 7:3, 5:5], [9:1, 6:4, 3:7], [9:1, 5:5, 1:9], [8:2, 6:4, 4:6], [8:2, 7:3, 3:7]. For example, [9:1, 9:1, 9:1] represents average sequencing depths of 9× and 1× for each of the two strains, respectively at all three time points. In each repetition, every species was simulated according to a scenario randomly selected from the predefined scenarios. Paired HiSeq-2000 reads with the length of 100bp were generated from these two genomes using ART 2.5.8. At each time point, simulated reads of all 20 species were pooled together to get the microbial community data.

We subsequently processed the simulated dataset using four methods, ConStrains 1.0, MIDAS 1.3.0, StrainPhlAn 3.0, and LongStrain. Since StrainPhlAn and MIDAS did not tackle the non-dominant strains, only the results of the dominant strains were compared. The performances of three methods were evaluated by the recall and accuracy of the identified SNVs. Since our method and StrainPhlAn can flexibly select the reference genome, all SNVs were identified using the genomes in MIDAS database as the reference genomes. For MIDAS, the raw sequencing data were processed with default parameters at each time point and merged as a final result using its built-in module “merge_midas”. All sites with a dominant allele that was different from the reference genome were regarded as variants. For StrainPhlAn, consensus sequences were obtained at each time point and phylogenetic trees were constructed under “accurate” mode. As with MIDAS, SNVs identified by StrainPhlAn were also defined as genotypes different from the reference genome. However, StrainPhlAn cannot merge results from different time points and we therefore calculated the accuracies of StrainPhlAn at each time point separately.

To calculate the accuracy and recall of the identified SNVs, we had to get the true SNVs of the strains relative to the reference genome. Therefore, we generated reads from the original genomes of the strains to a high depth of 100× using ART (Huang et al. 2012). These reads were mapped to the reference genomes in MIDAS database by Bowtie2 (Langmead and Salzberg 2012) with default parameters. Then, SNVs were called by SAMtools 1.9 (Li et al. 2009) with the command “samtools mpileup -uf reference.fas sample.bam | bcftools call -c --output-type v -v”. These SNVs were regarded as the gold standard in the calculation of recall and accuracy. The proportion of SNVs identified by each method overlapping with the gold standard were calculated as the precision. The recall of each method was the proportion of variants in the gold standard that were correctly detected by that method.

For the comparison with ConStrains, the MAE and RMSE of proportion estimate were calculated as follows:

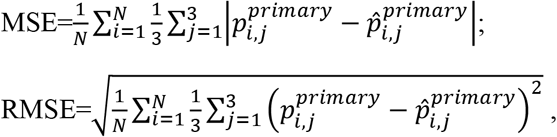

where *N* is the number of successful reports in 20 repetitions of the simulation, *i* is the index of repeat, *j* is the index of time point, and *p^primary^* is the proportion of the primary strain.

### Simulation study of single species

In this setting, we simulated each species separately as described in the simulation of the Gut20 community, but did not pool them together. For all 20 species, the raw sequencing reads were generated under two scenarios [9:1, 6:4, 3:7] and [8:2, 8:2, 8:2]. For the simulation of *B. breve*, we also conceived the scenarios with the same ratio at depths of 5× and 20×. To test the applicability of these methods under various scenarios, the raw reads of *B. breve* were also generated under the following scenarios: [6:4, 6:4, 6:4], [7:3, 7:3, 7:3], [8:2, 7:3, 3:7], [9:1, 7:3, 5:5]. The analysis and evaluation were performed in the same way as in the simulation study of the Gut20 community.

### Real data analysis

#### Analysis of a subset of TEDDY

In this study, we only focused on the subjects with T1D case-control status which consisted of 100 cases and 100 controls. We used LongStrain to screen 44 high-abundance species mentioned in two published TEDDY microbiome studies (Stewart et al. 2018; Vatanen et al. 2018).

#### Association between strain proportions/diversity and birth delivery mode

The log2-ratio of the proportions of the primary strain to the secondary strain at each time point was calculated by log_2_ 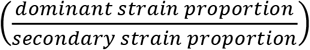. To reduce the noise, given each species, only time points with >3000 reads were retained and the log2-ratios were averaged over a sliding window of 2 months. To ensure an adequate sample size and include strain transitions, we focused on 26 species and used the samples from the first two years. We fitted the following linear mixed model to test whether the temporal trend of the log2-ratio was associated with the birth delivery mode.

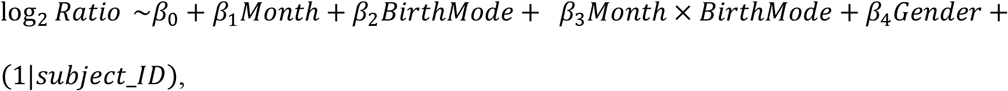

where “*BirthMode*” is the birth delivery mode (caesarian or vaginal) and “*subject_ID*” is a random effect. The gender was adjusted in this model and *p*-values were reported after FDR correction. Shannon’s index of strain proportion is calculated by 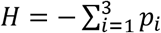 ln *p_i_*, where *p_i_* is the proportion of *i*^th^ strain at a time point. Similarly, we fitted the linear mixed model below to test the association between the temporal trend of strain diversity and the birth delivery mode.

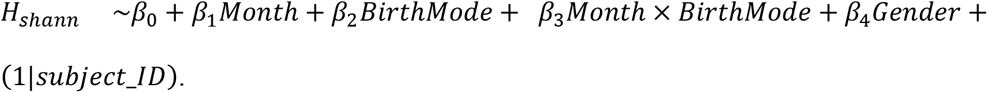

## Software availability

LongStrain is implemented in Python 3 and is freely available, along with step-by-step tutorials, at https://github.com/BoyanZhou/LongStrain.

## Data access

The raw sequencing files of the dataset from the TEDDY study is available in the database of Genotypes and Phenotypes (dbGaP; https://www.ncbi.nlm.nih.gov/gap/) with accession number phs001442.v1.p1. For the gastric intestinal metaplasia microbiome study, the metagenomic sequencing data is available in the dbGaP with accession number phs002566.v1.p1.

## Acknowledgments

This work was funded in part by the U.S. National Institutes of Health grants P20CA252728 and R01CA204113.

